# Full-waveform inversion imaging of the human brain

**DOI:** 10.1101/809707

**Authors:** Lluís Guasch, Oscar Calderón Agudo, Meng-Xing Tang, Parashkev Nachev, Michael Warner

**Author notes:** Contact: Lluís Guasch.

## Abstract

Magnetic resonance imaging and X-ray computed tomography provide the two principal methods available for imaging the brain at high spatial resolution, but these methods are not easily portable and cannot be applied safely to all patients. Ultrasound imaging is portable and universally safe, but existing modalities cannot image usefully inside the adult human skull. We use in-silico simulations to demonstrate that full-waveform inversion, a computational technique originally developed in geophysics, is able to generate accurate three-dimensional images of the brain with sub-millimetre resolution. This approach overcomes the familiar problems of conventional ultrasound neuroimaging by using: transcranial ultrasound that is not obscured by strong reflections from the skull, low frequencies that are readily transmitted with good signal-to-noise ratio, an accurate wave equation that properly accounts for the physics of wave propagation, and an accurate model of the skull that compensates properly for wavefront distortion. Laboratory ultrasound data, using *ex-vivo* human skulls, demonstrate that our computational experiments mimic the penetration and signal-to-noise ratios expected in clinical applications. This form of non-invasive neuroimaging has the potential for the rapid diagnosis of stroke and head trauma, and for the provision of routine monitoring of a wide range of neurological conditions.

## INTRODUCTION

No universally applicable means of imaging the living human brain at high anatomical resolution exists. The modality with the best spatial resolution and tissue contrast, magnetic resonance imaging (MRI), is contraindicated where the presence of magnetic foreign bodies cannot be excluded, and is impractical with claustrophobic, uncooperative or severely obese patients. Its nearest rival, X-ray computed tomography (CT), involves exposure to harmful ionizing radiation. Both require large, expensive, immobile, high-power instruments that are near-impossible to deploy outside specialized environments. The clinical consequences of this are high symptom-to-image times, long inter-scan intervals during serial imaging, and constraints on the range of patients that can be imaged successfully.

Pre-eminent amongst the many neurological disorders where patient outcomes are degraded by these restrictions, is stroke: the second most common cause of death worldwide, and the dominant cause of acquired adult neurological disability [1]. Treatment decisions are here critically guided by neuroimaging, ideally performed immediately after symptom onset. Delays of the order of minutes have substantial impact on outcomes, yet the necessity to treat patients only after transport to hospital routinely introduces delays of an hour or more [2]. Accelerating the treatment of stroke by enabling neuroimaging and treatment to be performed at the point of first contact would thus have large population-level impacts on survival and disability. Analogous arguments can be made for improved rapid medical imaging in head trauma, and in routine intraoperative, post-operative and preventative neurological monitoring, with the potential to impact large numbers of patients worldwide.

We provide *in-silico* proof-of-principle, supported by *ex-vivo* laboratory measurements, that the combination of transmitted transcranial ultrasound tomography with a computationally intensive technique originally developed to image the interior of the Earth, can address these clinical needs by providing portable three-dimensional (3D) quantitative imaging that is less-expensive, faster and more easily applicable than MRI, and that is safer and has better soft-tissue contrast than CT. This approach results in a three-dimensional, sub-millimetre resolution, quantitative model of acoustic wave speed within the brain and surrounding tissue, that is capable of distinguishing most of the structures and pathologies to which MRI is sensitive. The combined findings of our *in-silico* and *ex-vivo* experiments demonstrate that recording transcranial ultrasound data is feasible, recorded signal-to-noise levels are sufficient, the data contain the information required to reconstruct brain properties, and full-waveform inversion can extract those properties at high resolution.

Conventional medical ultrasound is fast, safe, portable and cheap, but is unable to image the adult human brain at high-resolution within the skull; the main reasons for this are well understood [3–6]:

1. In both conventional pulse-echo B-mode sonography [7] and time-of-flight ultrasound computed tomography [8], high-frequencies are required in order to obtain high spatial resolution. Scattering and anelastic losses occur within the skull and the brain, and these increase with frequency. At the frequencies used by conventional ultrasound modalities, these signal losses prevent successful imaging of intracranial soft tissue.
2. The contrast in wave speed between the skull and soft tissues, and between the skull and air-and-fluid-filled cavities within it, produces significant refraction, diffraction and reverberation of ultrasound energy as it is transmitted through the skull. This significantly distorts and complicates the consequent wavefront, leading to strong aberrations in both phase and amplitude, and to significant spatial and directional variation in the waveform of the transmitted pulse. It is not currently possible to correct for these effects with sufficient accuracy using conventional modalities.
3. In pulse-echo sonography, back-scattered reflections are used to generate the image. The bones of the skull differ significantly in wave speed and density from those of surrounding soft tissue. Consequently, the skull generates strong reflections and multiple scattering, and these high-amplitude signals overlie, interfere with and obscure the much-weaker reflections produced by the small impedance contrasts that occur between tissue types within the soft tissues of the brain, leading to low signal and high source-generated noise in intracranial pulse-echo images.
4. Time-of-flight tomography uses a short-wavelength approximation, basing its analysis on the simplified physics of ray-theory in which the effects of transmission through a heterogeneous medium are represented by a simple change in travel time. For a finite-wavelength wave transmitted through a medium that is heterogeneous on many scales, such delay times are only sensitive to the properties of the medium averaged over the dimensions of the first Fresnel-zone [9]. Consequently, time-of-flight tomography is unable to resolve structure below this scale, and so lacks acceptable resolution at the low frequencies that can be recorded using transcranial ultrasound.

One possible way around these problems is to use natural openings in the skull as acoustic windows, but this approach severely reduces illumination [5]; it is typically limited to neonates through an open fontanelle [10]. In principle, it is also possible to remove, or thin, portions of the skull in order to record data without strong bone reflections. This method has produced promising results in rodents, generating functional ultrasound images that can capture transient changes in blood volume related to brain activity [11, 12], but this invasive approach has obvious limitations in clinical practice.

In this paper, we present the neurological application of full-waveform inversion (FWI) [13], an imaging method first applied widely in geophysics [14]. FWI is a computationally intensive technique that has been developed to a high level of sophistication by the petroleum industry to image hydrocarbon reservoirs within the Earth [15, 16]. The spatial resolution that can be obtained using this technique is much greater than that of time-of-flight tomography. FWI achieves this improved resolution through a combination of characteristics [17], of which the most important is that it uses a more-complete description of the physics of wave propagation in heterogeneous media that takes proper account of the finite wavelength of transmitted waves. This description, which involves the full numerical solution of the wave equation, is able to model accurately the effects of sub-Fresnel-zone heterogeneity and multiple scattering on the wavefield. FWI combines this more-accurate description of the physics with an appropriate non-linear inversion scheme, and a suitable data acquisition geometry, so that it is able to recover fine-scale heterogeneity throughout the model.

Fig. 1 outlines the geometry of the method. Low-frequency ultrasound data are recorded at all available azimuths by surrounding the head with ultrasound transducers in three dimensions. Every transducer acts, in turn, as a source of ultrasound energy, and this energy is recorded by every other transducer. FWI uses predominantly transcranial transmitted energy recorded on the side of the head opposite to the source transducer, but it also extracts information from all other parts of the recorded wavefield including reflections, diffractions, multiple scattering and guided waves that arrive at any angle at any of the transducers. Unlike conventional ultrasound imaging, FWI does not use focused transducers, focusing arrays or any type of beam forming, either in the experimental configuration or in the computer subsequently.

**Figure 1.**
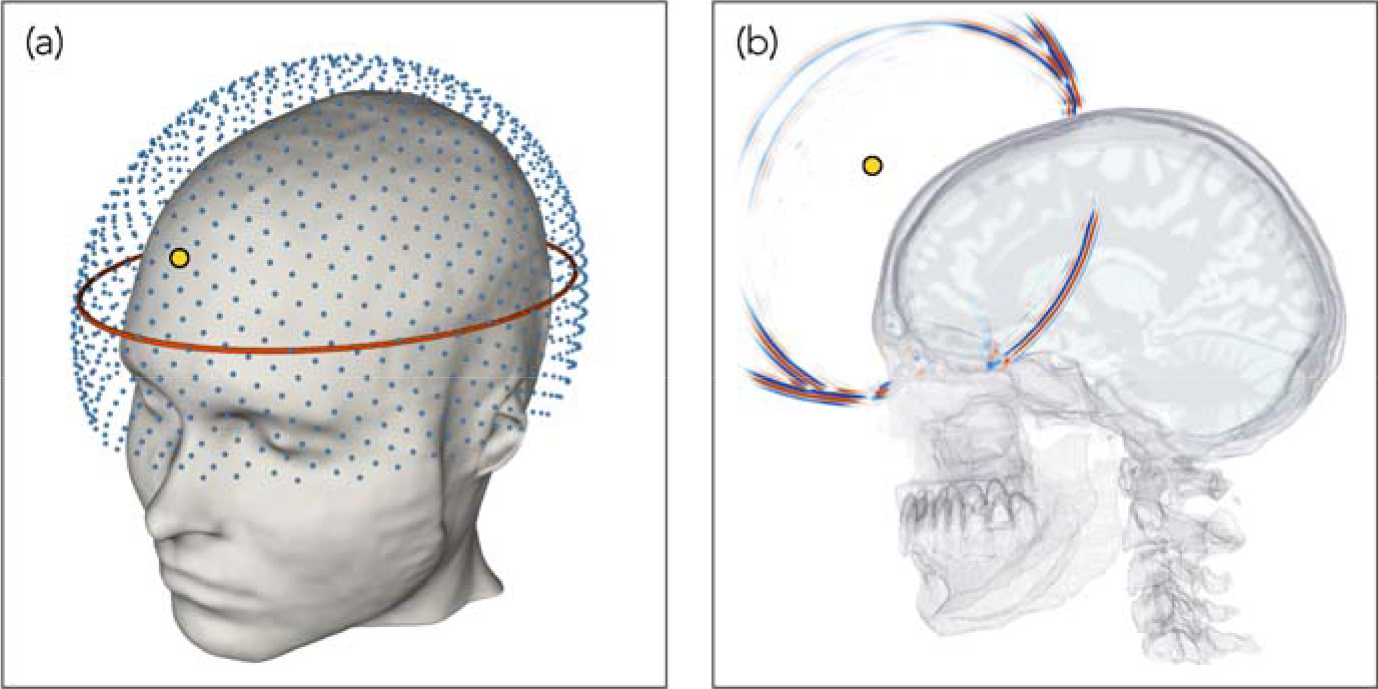
Experimental geometry. (**a**) Three-dimensional array of transducers used for data generation and subsequent inversion. Each transducer acts as both a source and a receiver. The red ellipse shows the location of the two-dimensional array used to generate the data for Figs. 2 & 4. (**b**) A snapshot in time of the wavefield generated by a source transducer located at the position indicated by the small yellow circle, computed via numerical solution of the 3D acoustic wave equation. The wavefield is dominated by strong reflections from the skull, and by intracranial transmitted energy travelling across the brain; Supplementary Movie 1 shows the full wavefield propagating in time.

The paper is organized as follows: We explore our proposed methodology using *in-silico* simulations, and present *ex-vivo* laboratory results that support our assumptions. We begin by demonstrating the improvement in resolution provided by FWI in even the simplest case when the model is two-dimensional and the skull is absent. We follow this by exploring what FWI is able to achieve for the intact adult human head in three dimensions; this result demonstrates the resolution and tissue contrast potentially achievable in a clinical setting. We follow this by demonstrating a practical method for building a starting model, and demonstrate the importance of full 3D data acquisition and inversion. We present laboratory results using an *ex-vivo* human skull to demonstrate that good signal penetration and high signal-to-noise levels are readily achievable by transcranial ultrasound. We provide an example of the clinical relevance of our approach by demonstrating the accurate recovery of an intracranial haemorrhage, and discuss clinical applications to stroke and other pathologies. We conclude with an outline of our methodology, and in the supplementary information we explore the consequences of potential imperfections in real-world data.

## RESULTS

### Resolution in the absence of the skull

Ray-based time-of-flight tomography and wave-equation-based FWI both represent forms of transmission tomography. Fig. 2 demonstrates the difference between these two techniques using a simple two-dimensional model of the naked brain without the complicating effects of the skull. Using the model from Fig. 2a, and solving a numerical wave equation, a synthetic dataset was generated for transducers located around the brain. Using the homogeneous starting model shown in Fig. 2b, this dataset was inverted using both time-of-flight tomography and FWI, to recover the models shown in Figs. 2c and d. Time-of-flight tomography seeks to find the best-fitting model by using geometric ray theory to predict delay times for every source-receiver pair in the dataset, whereas FWI seeks to solve the same problem by using the wave equation to predict the detailed variation of acoustic pressure with time recoded at every receiver for every source.

**Figure 2.**
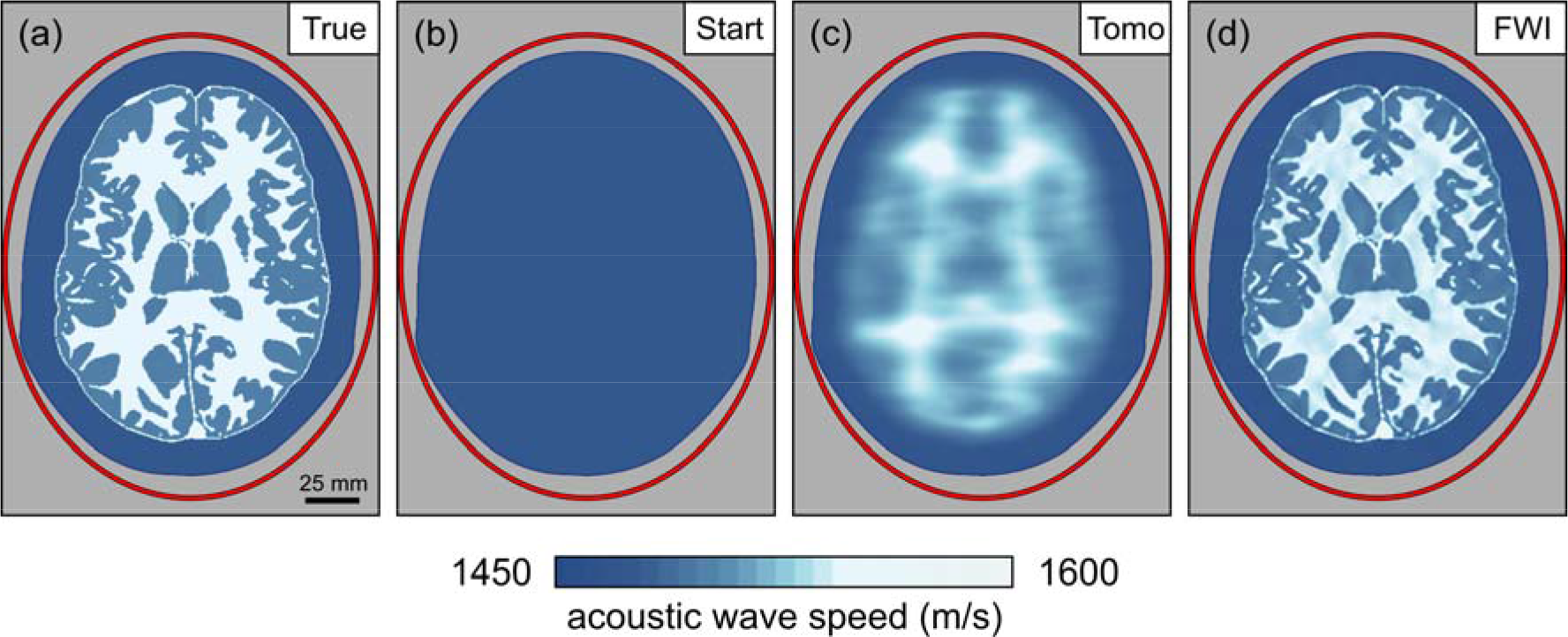
Inversion of data from a brain outside the skull. (**a**) A two-dimensional model of acoustic-wave speed in the naked brain without the skull. The red ellipse shows the transducer positions. (**b**) Homogeneous model used to begin inversion. (**c**) Result of ultrasound computer tomography. The resultant model is accurate but has poor spatial resolution. (**d**) Result of ultrasound full-waveform inversion. The resultant model is now both accurate and spatially well resolved.

For this numerical experiment, the shortest wavelength of the insonifying signal was less than 2 mm, and the minimum diameter of the Fresnel zone for signals that travelled across the model was greater than 20 mm. Well-established theory [9] and numerical experiments [18] show that the maximum spatial resolution that can be achieved, in the far field, using ray-based time-of-flight tomography is of the order of the diameter of the first Fresnel-zone, whereas for wave-equation-based transmission tomographic methods the maximum achievable resolution is of the order of half a wavelength [14, 19]. Thus, we would expect that the FWI model would be about 20 times better resolved in linear dimensions than the time-of-flight model. Fig. 2 illustrates this behaviour directly. Both methods recover models that are accurate in their locally averaged properties, but the time-of-flight model has only centimetre-scale spatial resolution whereas the FWI model has millimetre resolution. Note that, in this simple example, the difference in resolution between the two techniques is not related to the presence of the skull, nor to differences in the optimization scheme – both methods used non-linear least-squares inversion applied to the same input data.

In the absence of the skull, conventional high-frequency pulse-echo sonography would of course be able to recover an accurate image of the naked brain. However, when the skull interposes, pulse echo will fail to image the brain inside the skull because brain reflections are then significantly distorted by the skull, and signal loses at typical pulse-echo frequencies are large. Similarly, time-of-flight tomography for the intact human skull will fail because, at the low frequencies that can be transmitted across the head with acceptable signal-to-noise ratios, spatial resolution is insufficient. FWI does not suffer from either of these problems; it properly accounts for the distorting effects of the skull, and it achieves good spatial resolution at the low frequencies that can be recorded after transmission through the skull.

### Three-dimensional full-waveform imaging through the skull

FWI has obvious advantages for brain imaging; it does though have two complications of its own: the computational effort required to extract the image from the data in three dimensions is significant, and the method requires a reasonably good starting model in order to proceed to the correct final model.

The former requirement has, until recently, limited the applicability of medical FWI to problems that can be usefully solved in two dimensions [20], and the skull is not even approximately two-dimensional. The advent of large parallel multi-core multi-node compute clusters, of on-demand parallel cloud computing, and of large-memory GPU systems, coupled with improved FWI software and the use of pre-trained supervised deep-learning to accelerate the process, are reducing the computational demands of this method; runtimes and costs continue to reduce year-on-year.

The requirement for a good starting model is straightforward for the soft tissue of the brain where a homogeneous starting model is adequate. For the bones of the skull, either an accurate skull model must be obtained *a priori*, or a version of FWI must be employed that is able to build such a skull model from the observed data. In this section we assume that the skull model is known, and in the following section we demonstrate a method of building such a model during FWI.

Fig. 3 shows transverse, sagittal and coronal sections through a three-dimensional target model of wave speed, a starting model containing the true skull but otherwise homogeneous, and the model reconstructed using FWI applied to sub-MHz ultrasound data generated by the target model. Supplementary Movies 2, 3 and 4 show the true, starting and reconstructed models in three dimensions. The colour scale shown in Fig. 3 is designed to highlight heterogeneity within both soft and hard tissues.

**Figure 3.**
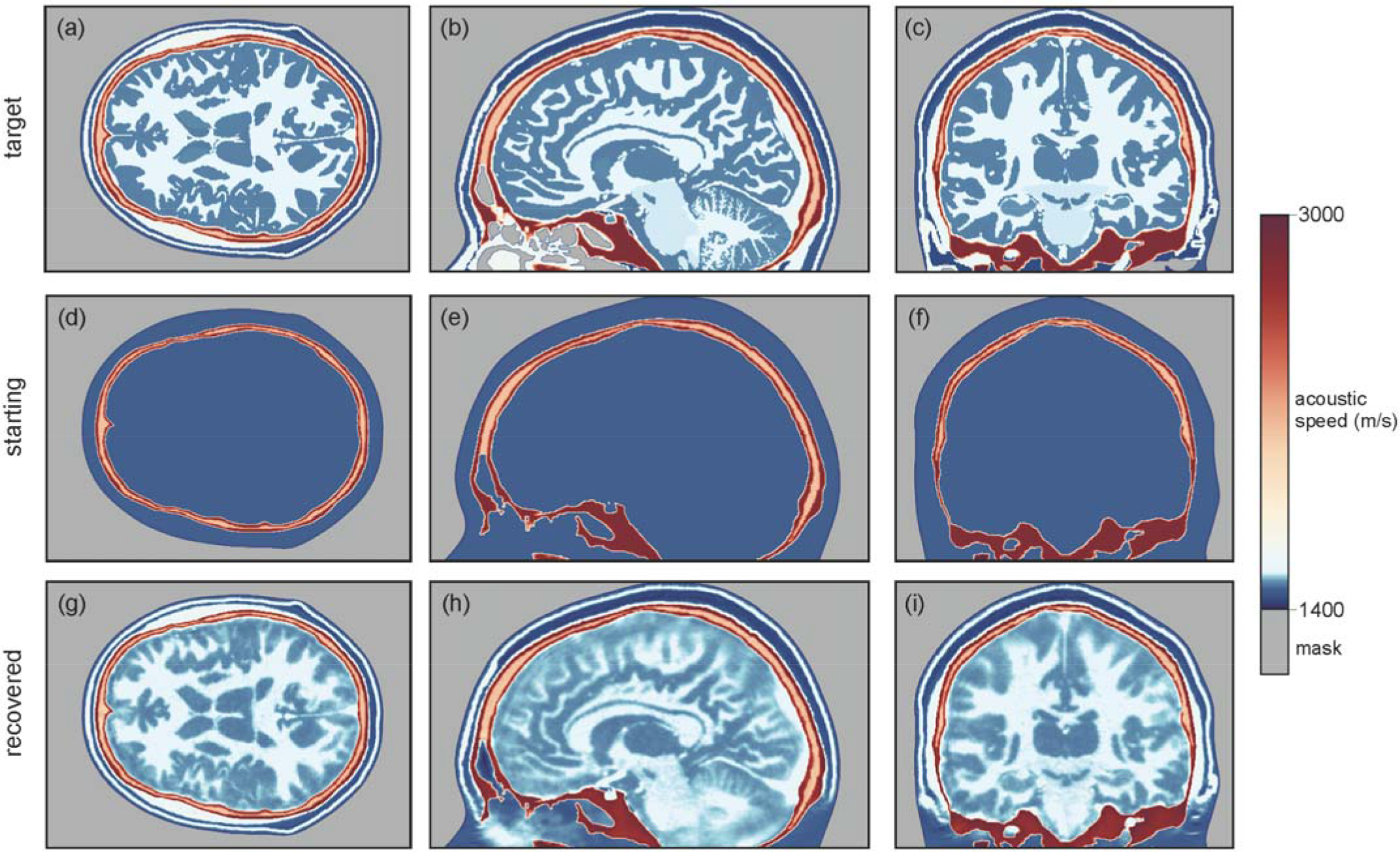
Models of acoustic wave speed. Transverse (left), sagittal (centre) and coronal (right) sections through the true (top), starting (middle) and recovered (bottom) models. Both the wavefield modelling and waveform inversion are performed in three dimensions. The starting model includes the true model of the skull, but is otherwise homogeneous.

The bones of most of the upper cranium are multi-layered, containing the inner and outer tables of denser cortical bone with a high wave speed, surrounding the diploë, which is formed of cancellous bone with lower density and wave speed. This structure, together with the large contrast in properties between the skull and its surrounding soft tissues, provides the principal mechanism for transcranial signal attenuation, with anelastic absorption and elastic mode-conversions playing a less significant role [21–23]. The model of the skull used in this study included all cavities, foramina and other structural complications that are present in the adult human head, and that are capable of being captured on the 500 micron grid that we used to represent the model.

The model recovered by full-waveform inversion, Fig. 3g to i, is in good agreement with the true model, Fig. 3a to c, for both extracranial and intracranial soft tissues. Inside the skull, FWI is able to generate an accurate and detailed image: grey and white matter match the target tissue properties accurately, both in absolute wave speed and in structure, with sufficient resolution to allow direct identification of cortical folds. Deeper structures such as the corpus callosum, the thalamus, the basal ganglia, and the ventricular system are recovered well. Parts of the venous sinuses have a thickness of 0.8 mm in the true model, as do larger vessels within the brain, and these are recovered in the reconstructed image demonstrating that we are able to achieve sub-millimetre resolution of the brain and its vascular system using only relatively low frequencies lying below 1 MHz. Parts of the cerebellum and the pons lie inferior to the lowest transducer positions in our numerical experiment, but it is still possible to extract sufficient information from the data to image both bodies, although there is a decrease in resolution as illumination is progressively lost in the area close to the base of the skull.

### Building the skull model

FWI is a local optimization algorithm that requires an initial model that lies within the basin of attraction of the global solution [14]. The variation in soft-tissue acoustic wave speed is around ±7%, which has values between about 1400 ms^−1^ for fat and 1600 ms^−1^ for muscle tissue and cartilage [24]. At the frequencies that we use for FWI, such relatively small perturbations are readily retrievable starting from a homogeneous model having a wave speed similar to that of water at about 1500 ms^−1^. This is the reason why FWI applied, for example, to breast imaging has been immediately successful [20, 25]. In contrast, the variation in wave speed for hard tissue in the cranium is larger at around ±14%, with values between about 2100 ms^−1^ for cancellous bone and 2800 ms^−1^ for cortical bone [21–23]; the mandible and the vertebrae have even higher wave speeds of around 3500 ms^−1^ [24]. These high values are far removed from that of water. Consequently, recovery of the full model of the head, including the bones of the skull, will require a more-sophisticated approach.

Fig. 4a shows the failure of an attempt to recover a model of the head using conventional FWI beginning from a purely homogeneous starting model. This should be compared with Fig. 3g which shows the analogous result obtained when the starting model contains a model of the skull. One solution to building an adequate starting model for the skull would be to extend the frequency spectrum of the ultrasound source to include even lower frequencies, mimicking the approach commonly employed in geophysics [14]. Despite the ease and elegance of such a solution, we have not explored that approach here because currently available ultrasound transducers are not sufficiently broadband to allow its practical implementation within a single device. Instead, we demonstrate that a more-advanced form of FWI, so-called adaptive waveform inversion [26], is able to build the skull model, from a homogeneous starting model, using only the range of frequencies that it is straightforward to generate.

**Figure 4.**
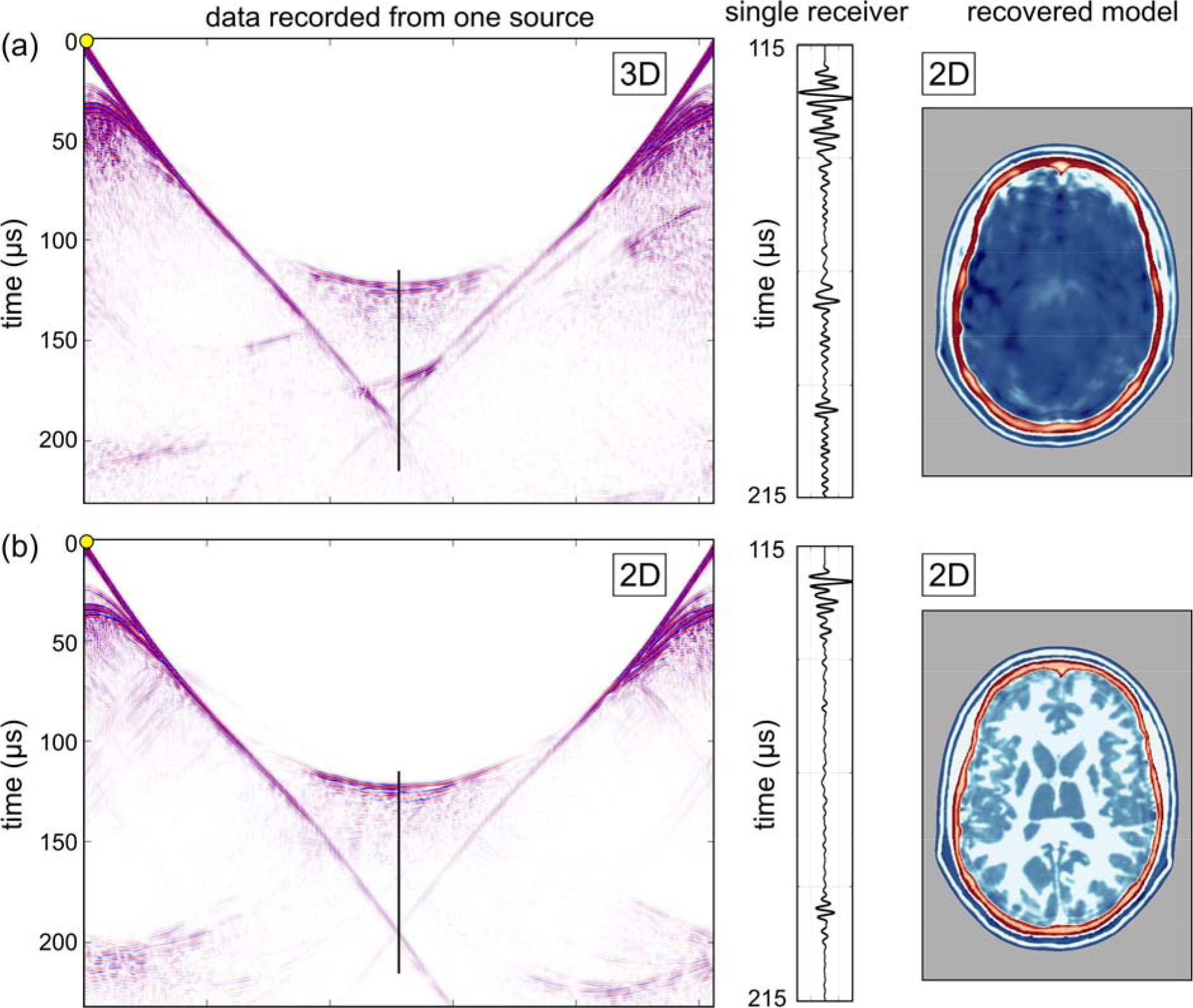
Model recovery using two-dimensional FWI. (**a**) 3D data inverted using 2D FWI. (**b**) 2D data inverted using 2D FWI. Left panels: show simulated data generated by a single source located at the yellow circle, as recorded on an elliptical array of 512 transducers placed around the head. Centre panels: show data recorded by a single receiver located opposite the source; the position of the data shown is indicated by the blue line. Right panels: show models recovered using purely two-dimensional FWI. The colour scale is as shown in Fig. 3.

Fig. 4b shows the result of applying adaptive waveform inversion, starting from the same homogeneous model as was used to generate Fig. 4a. Now the attempt at recovering a reasonable starting model for the skull purely from the data succeeds even though no *a-priori* model of the skull is assumed. Adaptive waveform inversion has immunity to cycle skipping, which is otherwise a common problem for conventional FWI [27], as well as an increased ability to recover sound-speed information from strong reflections such as those generated by the bones of the skull. We suspect that both of these characteristics may play a role in explaining the improvement of Fig. 4b over Fig. 4a. By analogy with the problem of imaging below high-contrast salt boundaries in geophysics [28], the recovered model of the skull would likely be further improved by the addition of total-variation constraints applied during adaptive waveform inversion, leading to a sharpening of the boundaries of the skull. The model in Fig 4b is now suitable for segmentation to extract the skull model, which can then be inserted into a homogeneous model, allowing the inversion to continue from this starting model using conventional FWI.

In practice, we suspect that such solutions may not actually be required in a clinical setting, and we have not pursued them further here. Large datasets consisting of many tens of thousands of MRI and X-ray CT images of adult human heads are becoming available [29], and ultrasound FWI datasets, both experimental and simulated from the other datasets, will also likely become available over time as transcranial ultrasound FWI develops. The pattern-recognition abilities of modern machine learning are already making significant impact in medical imaging and analogous areas [30]. It seems probable therefore that a suitably pre-trained deep neural network will be able to categorize raw reflection ultrasound observations against a suite of standard datasets to recover rapidly a parameterized model of the skull that will provide a high-quality starting model for FWI. This model would then be subsequently refined during conventional FWI of transcranial transmitted ultrasound to obtain the final quantitative image of both skull and brain.

### Importance of three dimensions

Most three-dimensional medical imaging analyses data initially in two-dimensions in order to produce a stack of planes that are combined to form a final 3D image volume. There would be advantages in applying this approach to ultrasound FWI: the computational cost of inverting many 2D slices is lower than that of true 3D inversion, and 2D acquisition systems are simpler to design, build and operate. However, the structural complexities of the skull, and large contrast with soft tissue, both act to distort the wavefronts by refracting and scattering energy out of the 2D plane.

Fig. 5a illustrates the detrimental effects of inverting three-dimensional data in only two dimensions. Here, the data being inverted are a dense two-dimensional subset of the full three-dimensional data used to generate the results shown in Fig. 3. In Fig. 5a, the data to be inverted have been generated by a 3D wave equation applied to a 3D model, but the inversion assumes only a 2D model and uses a 2D wave equation. The inversion is therefore unable to explain energy that has been refracted, reflected, scattered or guided out of the 2D plane. The model recovered in this case is neither accurate nor useful.

**Figure 5.**
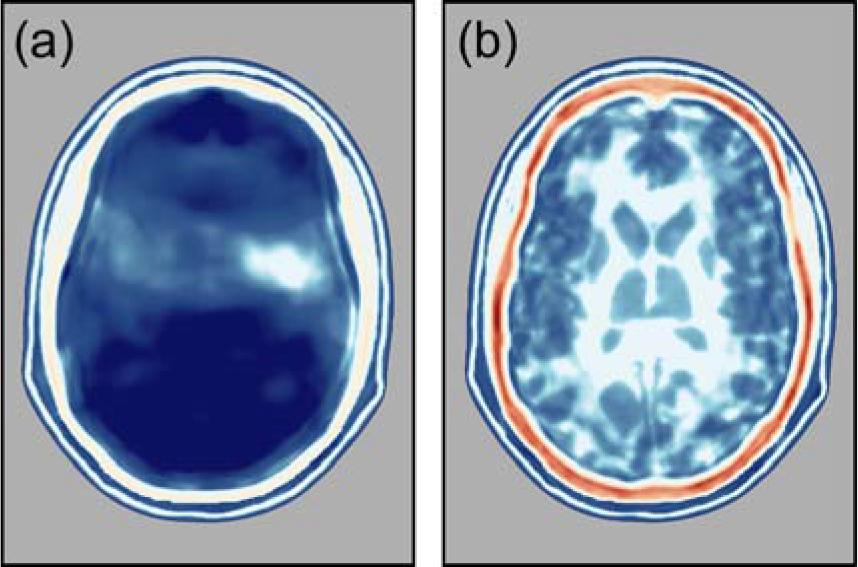
Inverting from a homogeneous starting model. (**a**) Model recovered using conventional FWI. (**b**) Model recovered using adaptive waveform inversion. The colour scale is as shown in Fig. 3.

Fig. 5b shows the equivalent experiment conducted purely in 2D; in this second case, both the initial data generation and the inversion are two-dimensional. The 2D inversion of 2D data recovers a model that is as accurate as that recovered by 3D inversion of 3D data. Comparing the data and waveforms in Figs. 5a and b demonstrates why 3D FWI of 3D data, and 2D FWI of 2D data both succeed, whereas 2D FWI of 3D data fails entirely. The 2D and 3D datasets show major differences, and it is evident that there must be significant out-of-plane energy present in the 3D data, and this cannot be explained adequately during 2D FWI. Since three-dimensional effects will always be present in real data, successful imaging of the brain using transmission FWI will always require 3D data acquisition and 3D inversion in order to correct properly for the three-dimensional distortion of the wavefield produced by the bones of the skull.

### Signal-to-noise ratios in ex-vivo experiments

At the sub-MHz frequencies that were used in the numerical experiments, transmission losses in soft tissue are small [24, 31–36], but scattering and anelastic losses in the skull can be important [21–23]. To test the significance of these losses, we conducted an *ex-vivo* experiment in the laboratory, immersing a real human skull in water, recording transcranial ultrasound using the same bandwidth and source amplitude as was used in the numerical simulations. The data were recorded, and are displayed in Fig. 6, un-stacked and un-processed. That is, a single source was triggered once, and the raw recorded data are displayed for each receiver without beam-forming, dynamic compression or other numerical manipulation. Absolute signal levels employed in this experiment were set to match those recommended by the British Medical Ultrasound Society for continuous adult transcranial diagnostic ultrasound [https://www.bmus.org/static/uploads/resources/BMUS-Safety-Guidelines-2009-revision-DETAILED.pdf].

**Figure 6.**
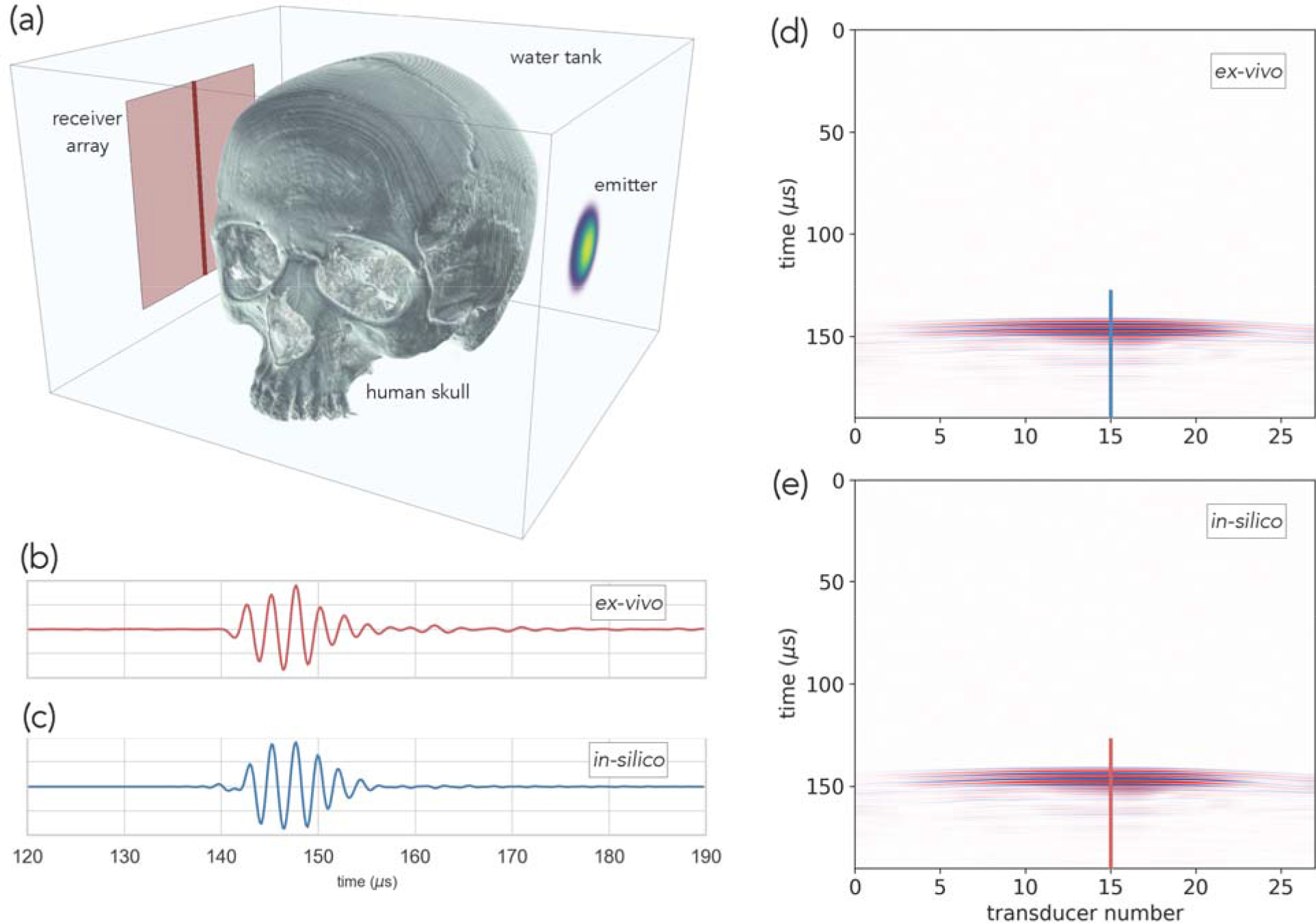
Ex-vivo and in-silico data after transmission across the head. (**a**) The geometry of the *ex-vivo* laboratory experiment. (**b**) Data recorded by the central *ex-vivo* transducer. (**c**) The equivalent *in-silico* data. (**d**) Laboratory data recorded on a finely sampled linear array. (**e**) The equivalent numerical data simulated in 3D. The physical skull and the numerical model are nominally the same, but differ in detail, and the numerical model does not include the effects of scattering by the physical transducers and their supporting hardware.

In the experiment, signals were transmitted across the cranium, passing through the bones of the skull twice, before being recorded. The source transducer was centred approximately on the squamous suture so that the source overlaps both the parietal and temporal bones. The principal signal losses within an *in-vivo* human head occur as the result of anelastic loss within bone, reflection at the inner and outer boundaries of the skull, and internal scattering within the bones of the skull. The principal wavefront distortions are produced by the large sound-speed contrast between the bones of the skull and their surrounding soft tissues, which have a sound speed close to that of water. This *ex-vivo* experiment is designed to capture all these features..

We repeated the same experiment *in silico*, using a model of the skull obtained by converting a high-intensity X-ray CT image-volume of the *ex-vivo* skull into acoustic sound speed. The conversion from X-ray attenuation to sound speed is not exact so that the *in-silico* data would not be expected to provide an exact match to the laboratory *ex-vivo* data. The real experiment will also contain signals scattered by the physical transducers and their supporting infrastructure; we did not attempt to duplicate these additional signals *in silico*.

Fig. 6 compares the two datasets. It shows that the timing, waveform, absolute amplitude and variation of amplitude with position and time, in the *ex-vivo* laboratory data, are well reproduced by the *in-silico* simulation. Most significantly, the noise level in the *ex-vivo* dataset observed in Fig. 6b is low, and low-frequency transcranial ultrasound, suitable for high-resolution FWI, penetrates the skull with only limited loss of signal intensity. In Supplementary Fig. 1, we show that FWI can tolerate much higher noise levels than are observed in the laboratory. Consequently, the signal penetration and signal-to-noise ratios that are likely to be achievable in practical applications will be more than sufficient for transcranial neuroimaging.

### Clinical application to stroke

The application of FWI to neuroimaging has the potential to improve diagnosis in a wide range of neurological pathologies; here we explore its potential to aid early treatment of stroke, a major cause of death and adult disability worldwide [1]. Stroke has two principal causes: ischemic stroke is most commonly caused by a blood clot obstructing blood supply to the brain, and haemorrhagic stroke is most commonly caused by bleeding within the brain parenchyma. When the blood supply to the brain is compromised, rapid intervention is required in order to restore circulatory integrity, halt and reverse tissue damage, and prevent and reduce morbidity, mortality and disability. While there are a number of early treatments available, including thrombolysis, mechanical thrombus extraction and similar interventions [37,38], their applicability is limited in practice by the requirement for accurate high-resolution brain imaging before these treatments can be deployed [2].

The indicated treatment for ischemic stroke is contraindicated for haemorrhagic stroke; brain imaging is therefore required to diagnose and separate these causes. The need for speed is paramount, but MRI is not portable and X-ray CT is barely so. Brain imaging then takes place not when paramedics first reach the patient, not within the ambulance, and not often within accident and emergency units; as a result, relatively few stroke patients receive a brain scan of any kind within the critical first hour, and even fewer receive high-quality MRI [2]. There is then a clear need for portable, fast, high-resolution, high-fidelity, 3D, brain imaging that can differentiate between ischemic and haemorrhagic stroke, and differentiate these from other pathologies that can mimic stroke. The development and clinical application of such a method would dramatically increase the survival rate and reduce the severity of subsequent disability by enabling much earlier treatment at the point of first patient contact.

To test the viability of this concept, we modified the target model to contain a haemorrhage, Fig. 7a to c. To build this model, we used physical properties for blood-infused soft tissue [39,40]. Using a homogeneous starting model for the brain, and the true model for the skull, we recovered the FWI images in Fig. 7d to f. Fig. 8 shows that the haemorrhage can be readily segmented from the three-dimensional model, both in the target and the FWI-recovered models. The target pathology is well recovered in the FWI image, and has good tissue contrast with other features of the brain. The boundaries and exact extent of the pathology are clear; and although it is not shown here, FWI is well able to produce time-lapsed images over a wide variety of time scales from seconds to hours. The spatial resolution of FWI is capable of detecting haemorrhage at all scales down to that of the originating vasculature.

**Figure 7.**
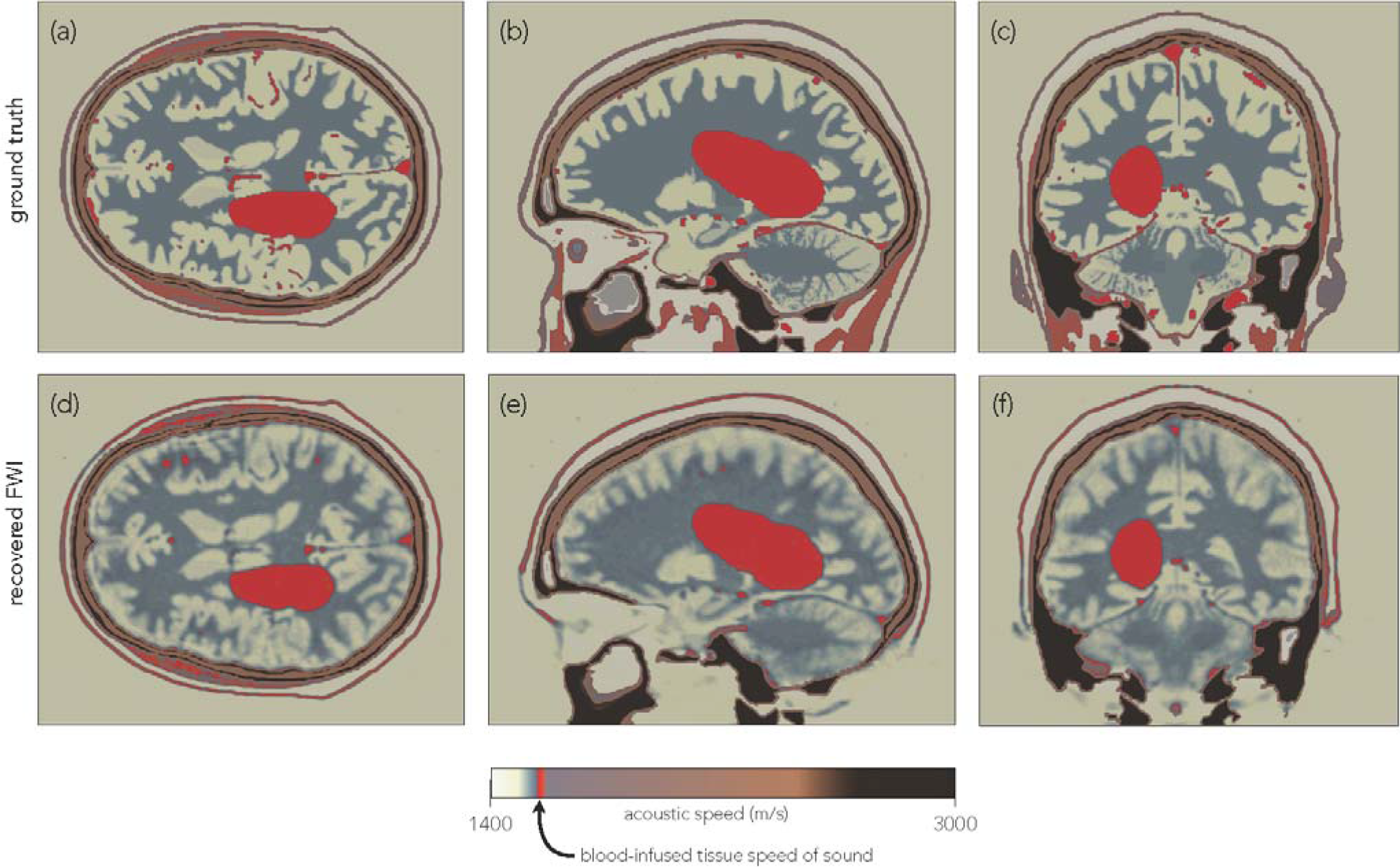
Recovery of a large haemorrhage by FWI. (**a** – **c**) Slices through a 3D wave-speed model perturbed by a large haemorrhage. (**d** – **e**) The same slices through the FWI. The haemorrhage is well recovered by full-waveform inversion at high resolution in all slices. The colour scale has been modified to highlight the haemorrhage.

**Figure 8.**
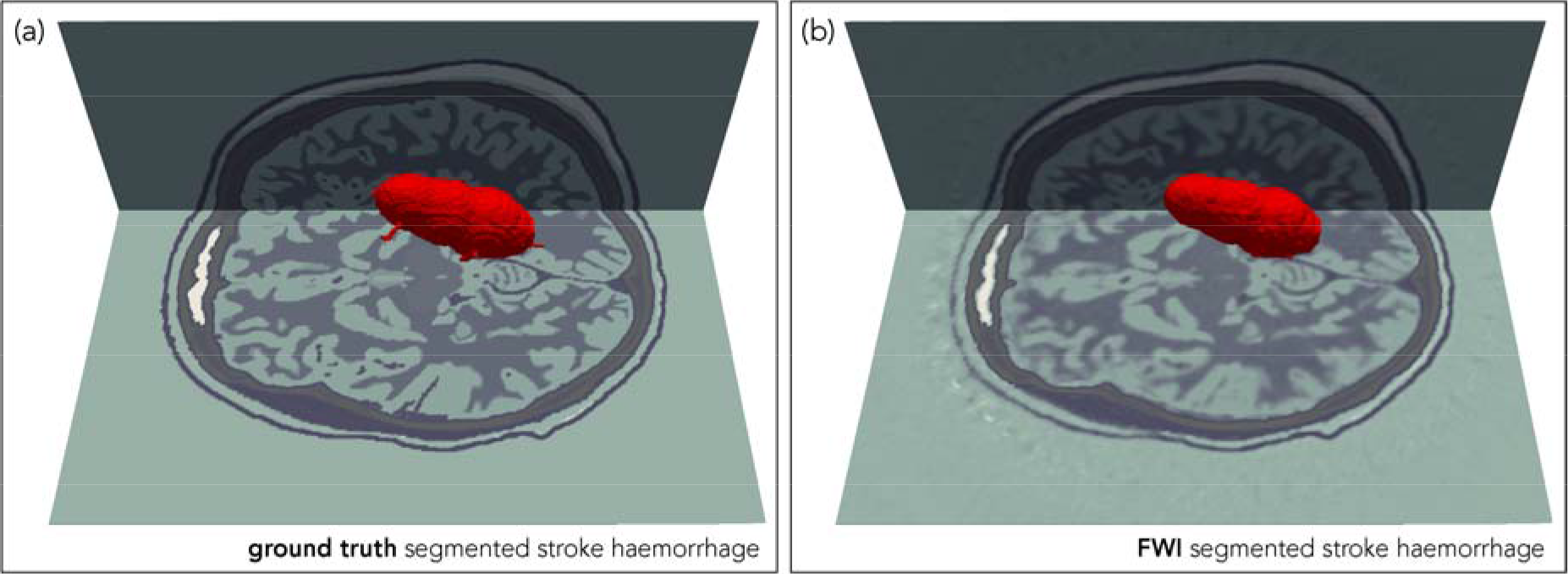
Segmented haemorrhage. **(a)** Haemorrhage auto-segmented from the true model. **(b)** Haemorrhage auto-segmented from the model recovered by full-waveform inversion.

## DISCUSSION

Both X-ray CT and MRI revolutionized medical imaging when they first appeared; three-dimensional transmission and reflection ultrasound tomography using full-waveform inversion has the same potential for impact across multiple disciplines, and has especial relevance for rapid diagnosis and treatment of stroke. Supplementary Fig.1 shows that the method is robust against the levels of noise that we observe in realistic *ex-vivo* laboratory experiments, and transcranial ultrasound signals have large amplitudes at the relatively low frequencies (< 1 MHz) that are sufficient for successful sub-millimetre resolution. The method overcomes the well-known limitations of conventional pulse-echo ultrasound imaging of the adult human brain, and the related limitations of conventional time-of-flight tomography.

It is necessary either to include some prior model of the skull within the starting model before attempting to recover the brain, or to use an advanced form of FWI such as adaptive waveform inversion that can converge towards the correct answer from a simpler starting model. In order to correctly account for and remove the distorting effects of the skull, it is essential to acquire and invert the data in three-dimensions, and unlike many other imaging techniques, it is not possible to reduce three-dimensional imaging merely to the sum total of a sequence of two-dimensional slices. In Supplementary Fig. 2, we show that it is not necessary to include density or anelastic absorption explicitly in the inversion in order to recover a good image, but it may be desirable to do so, both to improve the accuracy of the final image, and to obtain additional independent parameters to aid diagnosis.

The computational effort required for 3D FWI is considerable. The results shown in Fig. 3 require about thirty-two hours elapsed time to complete, running on a conventional cluster of 128, CPU-based, 24-core, compute nodes. Our target is to reduce this time to below ten minutes; this requires a speed up of about two-hundred times. The hardware that we used has a peak performance of about 60 tera-flop, so achieving the desired speed up requires hardware capable of operating usefully at a peak of about 12 peta-flop. Individual high-performance GPU-based servers are currently able to achieve speeds in excess of 1 peta-flop, so that a small array of these would in principle be capable of producing a final model in less than ten minutes. Assuming 2019 prices, amortization over three years, and full utilization of the hardware, the capital cost of the GPU-server hardware required to do this represents a few tens of dollars to invert a 3D transcranial dataset on a 500-micron grid. So, while the computational burden of FWI is high, the cost per patient is not high under appropriate circumstances.

The potential value of FWI imaging is three-fold. Most importantly it could improve outcomes in acute neurological disorders such as stroke and head trauma by enabling earlier intervention; the ultimate aim is diagnosis and treatment within minutes of first contact with paramedics. Second, the low cost, high safety, portability and high resolving power of the technology provides the ability to monitor the brains of patients continuously at the bedside allowing clinicians to intervene, for a range of pathologies, to prevent injury with the speed that the brain demands, acting in rapid response as if the brain image was a simple physiological variable such as blood pressure. And third, the technology can be deployed readily and safely, for prevention and diagnosis, in a wide range of situations where neuroimaging would be desirable but is currently unavailable – for example: within developing nations with limited health budgets, in remote locations, routinely at contact-sports events, within military deployments, or as part of disaster relief when local infrastructure is compromised.

## METHODS

### In-silico model

We used the MIDA 3D numerical model of the human head [41], at the original sample spacing of 500 microns, as the basis to build the sound-speed model used in the *in-silico* simulations. Physical properties within the model were derived from the geometry of the segmented model combined with values for acoustic sound speed, density and absorption for different tissue types from [21–24,31–36]; further details appear in Supplementary Table 1. Most minor tissue types within the model have unmeasured acoustic properties; in these cases, we estimated their values using small perturbations to the properties of other tissue types that appeared analogous in their other physical properties and composition.

The models used to generate the data for all figures except Supplementary Fig. 2 were purely acoustic. The model used for Supplementary Fig. 2 included anelastic absorption, and assumed a linear relationship between attenuation and signal frequency. At the relatively low frequencies used in these simulations, such a model of absorption provides a reasonable approximation to the properties of real tissue [31].

### In-silico modelling

The experimental geometry was restricted to accommodate the application of this technology realistically to human patients in a clinical setting, and therefore no transducers were positioned in front of the face or below the base of the head. Transducers were modelled assuming unfocused single elements. We used 1024 transducers that acted as both sources and receivers, generating just over a million source-receiver records of acoustic pressure, each record lasting 240 μs. The source waveform consisted of a three-cycle tone burst having a peak amplitude at a frequency of 400 kHz and a useful bandwidth for FWI extending from about 100 to about 850 kHz.

The synthetic data were generated by solving the three-dimensional, variable density, acoustic wave equation, explicitly in the time domain using, using a time-stepping finite-difference algorithm, with an optimised stencil that is nominally tenth-order in space and fourth-order in time. For the two-dimensional data shown in Fig. 4b, the analogous two-dimensional wave equation was solved in which the model, wavefield, sources and receivers do not vary perpendicular to a two-dimensional plane. For the anelastic data used to generate Supplementary Fig. 2c, the visco-acoustic wave equation was solved using the method described in [42].

### Full-waveform inversion

FWI is an imaging method that seeks to find a model that can numerically reproduce experimental data. It does this by solving a non-linear least-squares local optimization problem, modifying the model in order to minimize the misfit, *f*, defined as (half) the sum of the squares of the differences between the experimental data and an equivalent simulated dataset that is numerically generated using a numerical model of acoustic properties. Thus we seek to minimise:

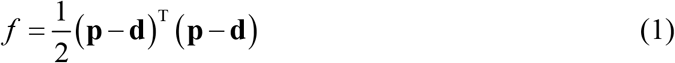

where **p** and **d** represent the predicted (numerically generated) and observed (experimental) data respectively organised as vectors that contain concatenated time-series of pressure variations at each recording location for every source location. In this *in-silico* study, the experimental data were themselves generated using a known ground-truth target model. At the initiation of FWI, a starting model **m**, representing an initial estimate of the target of interest and composed of many model parameters *m_i_*, is used to solve the wave equation and generate the predicted dataset **p**.

The computational cost of numerically solving the governing wave equation in 3D for many sources restricts computationally tractable solutions to iterated local gradient-descent methods. We solve the problem by seeking the direction of steepest descent, on the hyper-surface defined by *f*, which has as many dimensions as there are model parameters *m_i_*. For the 3D model shown in Fig. 3, this hyper-surface had about 10^8^ dimensions. This gradient-descent algorithm seeks to move from the starting model, by a sequence of small steps, successively downhill on this hyper-surface, to arrive close to the model that lies at the lowest point on the surface – this is the model that best predicts the observed data in a least-squares sense. Further details are provided in the Supplementary Information.

In a practical FWI algorithm, the direction of steepest descent is typically preconditioned in some way to speed convergence; here we used spatial preconditioning to compensate for illumination variation within the model [15]. We inverted the *in-silico* data over finite-frequency bands, starting at a dominant frequency of about 100 kHz, and moving successively to higher frequency to reach to a maximum dominant frequency of around 720 kHz. This multi-scale approach helps to ensure that the inversion does not become trapped at some local solution produced by inadequacies in the starting model. The highest frequencies present in the data have a half-wavelength in the brain of less than a millimetre, so that we would expect to be able to resolve sub-millimetre structure in the final recovered model.

### Time-of-flight tomography

The model shown in Fig. 2c was generated using time-of-flight tomography. This method is analogous to FWI, but with two significant differences: the data to be inverted consist of single numbers, one for each source-receiver pair, representing the time taken for energy to travel through the target model from source to receiver; and geometric ray-tracing rather than the full wave equation is used to calculate these travel times. This approach has the advantage that it is computationally more tractable than FWI, and so runs orders of magnitude more quickly; it is also much less likely than FWI to become trapped in local minima. But it has the disadvantage that it cannot recover structure in the model below the scale of the diameter of the first Fresnel zone. For a signal of wavelength *λ*, and a source-receiver separation of *x*, this diameter is approximately 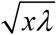, or about 17 mm for transcranial signals at 1 MHz, and this poor spatial resolution is apparent in Fig. 2c.

We solved the tomographic problem by minimising a misfit similar to that shown in equation (1), but where **p** and **d** were now vectors containing the travel-times rather than the raw observed acoustic pressure data. We address the problem as before by solving a non-linear least-squares local optimization problem, using gradient descent preconditioned by conjugate gradients. Unlike the more familiar X-ray CT, in ultrasound tomography changes to the model affect the path that energy follows from source to receiver, and it is necessary to include this non-linear effect into the inversion by iterating.

### Adaptive waveform inversion

Adaptive waveform inversion (AWI) is a form of FWI that has immunity to cycle skipping [26,27]. In conventional FWI, the algorithm seeks to drive the sample-by-sample difference between the predicted and observed data to zero. In contrast, in AWI the algorithm seeks to drive the ratio between the two datasets to unity. Both approaches aim to drive the predicted dataset towards the observed dataset, and consequently to drive the recovered model towards the true model. With perfect data, and perfect algorithms, both approaches will reach the same end point. However, when both methods are implemented using local gradient descent, they follow different paths through the space of possible models in their attempt to reach the true model. In these circumstances, FWI will tend to become trapped in local minima when the predicted data differ from the observed data by more than half a wave cycle, whereas AWI will not.

The ratio used in AWI is not generated sample-by-sample; rather it is a ratio formed frequency-by-frequency after temporal Fourier transform, and the ratio is formed separately for each source receiver pair. Since division in the frequency domain represents deconvolution in the time domain, the AWI algorithm effectively deconvolves one dataset by the other and then attempts to drive the result of that deconvolution towards a unit-amplitude delta function at zero temporal lag. The mathematical details are given on [26], and practical details are given in [27].

### Ex-vivo laboratory experiment

The laboratory experiment was performed by immersing a formalin-preserved *ex-vivo* human skull in water, generating an ultrasound pulse on one side of the head, and recording the resultant signals on the other. The skull retained some residual soft tissue, was stored dry, and was typically immersed for a few tens of minutes during ultrasound measurements. Sources and receivers were not in direct contact with the skull.

We used a single-element unfocused Olympus 500 kHz V381 19-mm ultrasound source transducer to generate a three-cycle tone burst centred on 400 kHz at the intensities commonly employed in conventional medical imaging [7]; this signal matches that used in the *in-silico* experiments. We recorded the transmitted signals from this emitter using an Olympus 500 kHz V301 25-mm transducer located on the opposite side of the skull at a perpendicular distance of 210 mm from the source. A planar receiver array was formed by moving the single receiving transducer successively in 4-mm steps to 729 positions to form a 27 × 27 array measuring 108 × 108 mm. The data were generated and recorded using a Verasonics Vantage 256, and the recorded data were low-pass filtered at 1 MHz.

## SUPPLEMENTARY INFORMATION

### Effect of noise on FWI

Our *in-silico* conclusions are only relevant to the real world if ultrasound signals, of the intensity required for medical imaging and with the bandwidth that we use here, can be recorded with sufficient signal-to-noise ratio after propagation across the head, traversing the bones of the skull twice in the process. Two questions are relevant: how sensitive is FWI to noise, and what level of signal-to-noise can be expected for transcranial ultrasound at the frequencies required? Below, we address the first of these questions; Fig. 6, in the main text, has answered the second.

Supplementary Fig. 1 shows the results of adding random noise to the *in-silico* data before subsequent inversion. Apart from the addition of noise, Supplementary Fig. 1 is exactly analogous to Fig. 4b. Comparison of these two figures demonstrates that models recovered with and without the addition of noise are similar; even the high level of noise applied here causes only modest degradation of the reconstructed image. FWI using transcranial ultrasound is clearly robust in the presence of high levels of incoherent noise. This robustness occurs because the formalism of FWI captures only those parts of the observed data that are capable of being reproduced by application of the wave equation to a model. Most forms of noise cannot be substantially reproduced in this way, and so, while noise may slow down convergence to the correct model, the level of noise in the final model is typically much lower than the level of noise in the data.

**Sup. Fig. 1.**
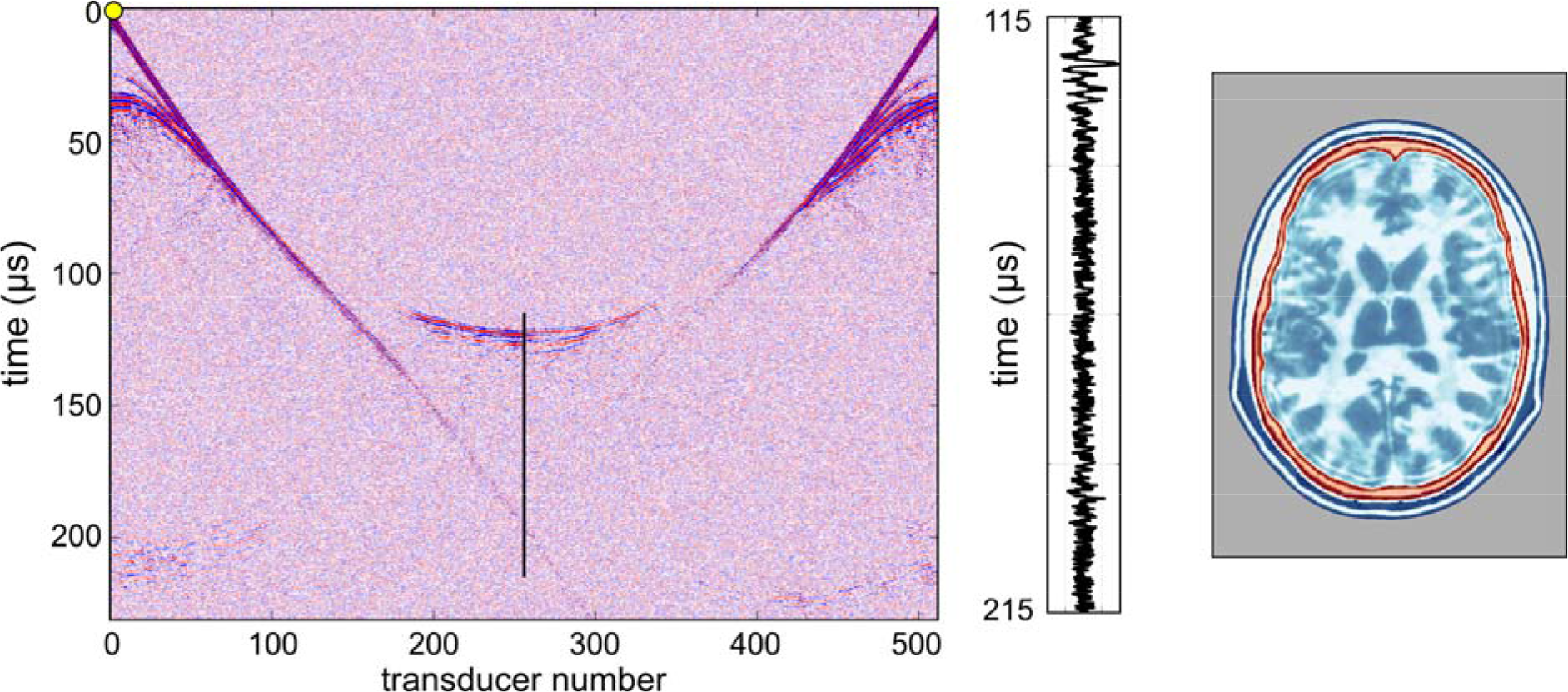
Model recovery using imperfect data. Fig. 4b presented 2D data inverted using 2D FWI. Here, we show the same data and the results of the same inversion, but after the addition of random noise to the raw data. The noise level here is much higher than the level of noise observed in the *ex-vivo* experiment shown in Fig. 6, and the recovery of the model shows only minor degradation as a consequence of the noise.

### Effect of absorption, density, anisotropy and elasticity

The modelling and inversion shown in Fig. 3 was performed without regard to anelastic absorption, density variation, anisotropy or elastic effects. Supplementary Fig. 2 shows the effects of ignoring both density and absorption during FWI. Absorption and density models for the head are shown in Supplementary Figs. 2a and 2b respectively. Here, absorption is displayed using quality factor *Q*, defined as 2*π* times the fraction of energy lost per wave cycle; lower *Q* values represent higher values of attenuation. Our model here assumes a linear relationship between attenuation and frequency, but it is straightforward to incorporate a more complicated relationship where appropriate.

**Sup. Fig. 2.**
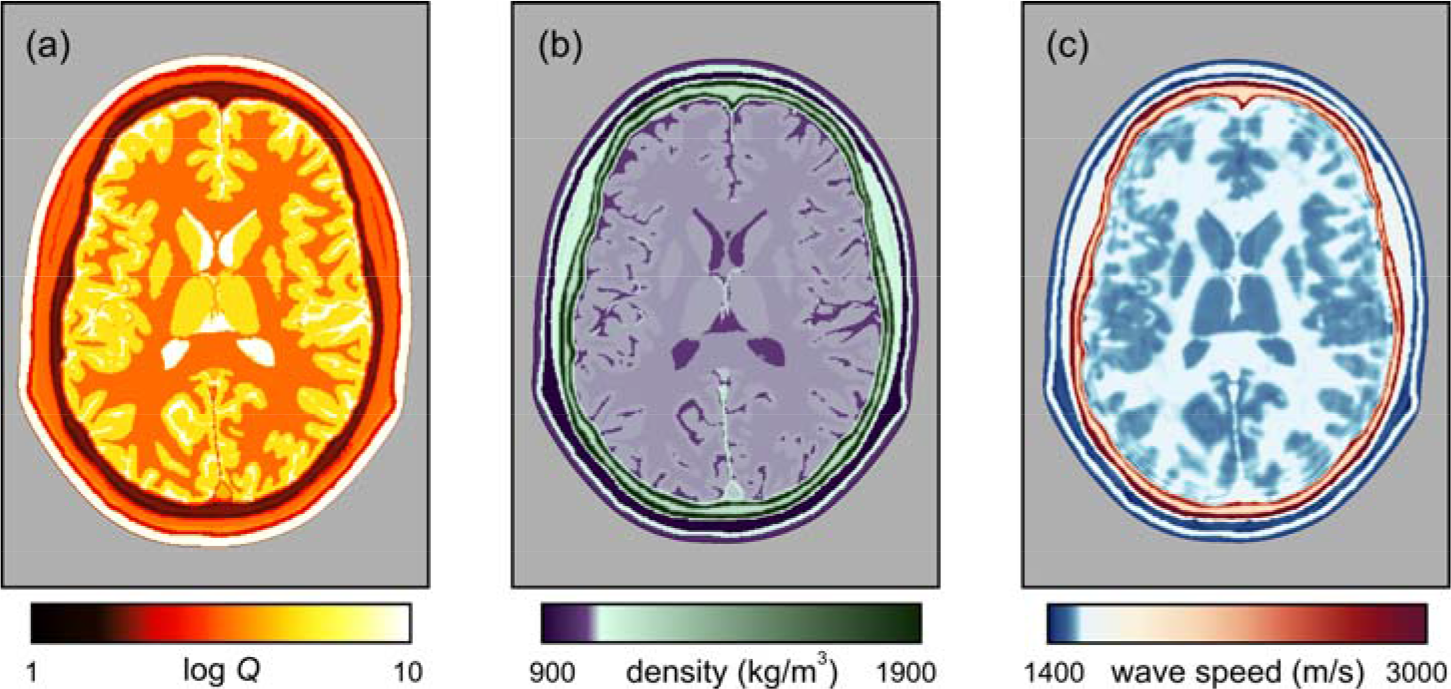
Models of the head. (a) Model of anelastic absorption. (b) Density model. (c) Sound speed recovered by FWI, using data generated including absorption and density, but assuming constant density and no attenuation during inversion. Simulations and inversions are both in 2D.

Supplementary Fig. 2c shows the model recovered by FWI assuming constant density and no attenuation when inverting simulated data that include both effects. A reasonable representation of the true wave-speed model is recovered. The principal effect of ignoring density variation is to ascribe all variation in signal amplitude to velocity variation, and this tends to exaggerate differences between locally heterogeneous regions. The principal effect of ignoring absorption and its associated velocity dispersion is to reduce the mean sound speed of the recovered model. Consequently, Supplementary Fig. 2c is slower on average than the true model, but its structure is nonetheless accurate. Ignoring both density and absorption effects during inversion is not significantly worse that ignoring either one alone. In clinical application, both of these effects can be included within the inversion – this is straightforward to achieve, but appears not to be necessary for the recovery of a simple well-resolved image. It is also possible to use FWI to invert explicitly for attenuation and other parameters [14], and some pathologies may be more readily diagnosed using such multi-parameter inversion than by use of acoustic wave speed alone.

Anisotropy in wave speed is routinely incorporated into FWI in geophysics [15] because it needs to consider crystalline, micro-fractured and finely layered materials. Other than perhaps in the skull, anisotropy in sound speed is unlikely to significantly affect ultrasound neuroimaging. Fully-anisotropic inversion for the skull is straightforward to incorporate into medical FWI, and most established 3D FWI codes will already have that capability incorporated.

Elastic mode-conversions in soft tissue are small; the skull however does have a significant shear modulus [21,22], and so can produce elastic effects in acoustic data. We have not yet investigated the quantitative importance of these phenomena in neuroimaging, but we do not expect them to be more significant than the effects shown in Supplementary Fig. 2c. Ultrasound sources and receivers in water do not generate or record shear waves, and elastic conversions are small close to normal incidence, which is the portion of the wavefield that is most important for FWI, both in transmission and reflection imaging. In geophysics, despite its potential importance, commercial FWI is seldom performed using the full-elastic wave equation, and the resultant models match well to in-situ direct measurements made within boreholes [14–16]. The Fullwave3D code used in this study is able to undertake anisotropic full-elastic 3D FWI, but its significant increased computational cost has not-often proven to be justified, at least in commercial geophysics.

In summary, acoustic, isotropic, constant-density, non-absorbing, three-dimensional inversion appears to be adequate for the generation of well-resolved detailed images of the brain. However, a more-complete account of the full physics during FWI, including absorption and anisotropy in bone, and elasticity in hard tissues, may help to provide a more quantitatively accurate model of physical properties, but at an increased computational cost which, at least initially, will translate into an increase in the total elapsed time required to compute the final model. Practical solutions are likely to involve simple fast acoustic inversion to form an initial image, followed optionally by more-accurate inversion using more-complete physics subsequently as circumstances require. In medical imaging, it is possible that additional physical properties recoverable by FWI will have diagnostic value advantages in special circumstances, as they do in both medical attenuation tomography [43] and in geophysics [44].

### FWI algorithm

Here, we continue the description of the FWI algorithm from equation (1) in the main text. In the account below, the quantities *f*, **p**, **A** and **u** each depend upon the assumed model **m**, whereas **d**, **s**, and **R** do not.

In order to find the direction of steepest descent, the derivative of the misfit with respect to the model parameters is found using the adjoint-state method [14]. The derivative of *f* with respect to each of the model parameters *m_i_* takes the form:

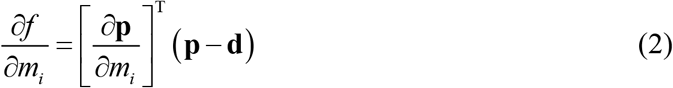

To compute the first derivative of the predicted data with respect to the model parameters *m_i_*, we start by writing the wave equation as a matrix-vector operation:

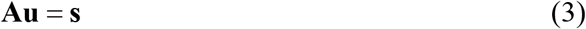

where **A** is the wave equation written in a suitable discrete form – here we used high-order finite differences to approximate the 3D anisotropic variable-density visco-acoustic wave equation, **u** is the pressure wavefield and **s** is the source. Differentiating this with respect to *m_i_*, and taking into account that both **A** and **u** depend upon the model parameter *m_i_* but that the source **s** does not, gives:

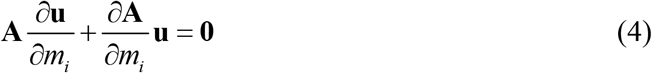

Assuming that **A** is invertible, which it must be if equation 3 has a unique solution, leads to the expression:

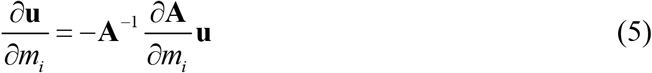

for the variation of the wavefield **u** with the model parameter *m_i_*. Now, the predicted data **p** are simply a subset of the full wavefield obtained at those locations where we happen to have placed receivers. Thus, we can use a restriction matrix **R** to extract the corresponding data as **p** = **Ru**, so that:

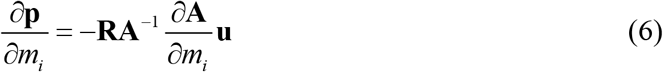

because, again, **R** does not depend on *m_i_*. The final expression for the gradient is then:

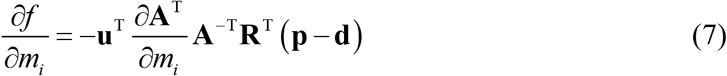

Reading from right to left, this expression implies the following sequence of steps to calculate the gradient:

1. compute the residual data (**p** − **d**) by solving the wave equation to generate **p** in the starting model,
2. inject the residual data into the model at the receiver positions,
3. solve the wave equation backwards in time using the injected residual data as a virtual source,
4. scale the resulting wavefield using the differential of the wave equation operator **A** with respect to the model parameters *m_i_*,
5. find the zero lag of the cross-correlation of this wavefield in time with the original forward wavefield **u** at every point in the model.

The result of applying these steps is a gradient vector oriented to point in the direction of maximum increase of *f* at the current position in the solution space. The negative of the gradient indicates the direction in which small changes to the model will create the largest decrease in the misfit *f*, so the model should be changed in this direction. Typically, the problem will be non-linear and non-convex, and therefore it requires that the model is updated iteratively, and that the starting point is within the basin of attraction of the global minimum. A more complete development, and further details are given in [13,14,17].

### Resolution of computer tomography and FWI

Spatial resolution in conventional pulse-echo ultrasound sonography typically depends upon pulse duration and beam width. Resolution in this context relates to the spatial discrimination of features seen in a reflection image around the location from which the ultrasound energy is reflected. In contrast, in transmission tomography, whether inverting the delay times used by conventional time-of-flight tomography, or the waveforms used by wave-equation FWI, it is the spatial discrimination of features by their effect on the transmission properties of the wave through the medium that is relevant. For time-of-flight tomography, in order to separate two adjacent features within a model, their effect on the recorded delay times must be distinct from the effect of a single feature that represents a smooth mixture between the two adjacent features. And for FWI, the separation of two features requires that their effect on the recorded waveforms must be distinct.

In this context, a Fresnel zone represents that region of a model through which energy can travel from a source to a receiver, arriving within the same half cycle. Such energy cannot be separated in terms of its delay time, and all such energy contributes to the same arrival. The first Fresnel zone corresponds to the region of the model through which energy travels that arrives within half a cycle the geometric ray arrival, which corresponds to the arrival time of an infinite-frequency wave travelling along an infinitely thin ray path. For a homogeneous model, the diameter of the first Fresnel zone at the midpoint along a path of length *x* is 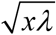, where *λ* is the wavelength of the signal. Since detail below the scale of the first Fresnel zone does not change observed delay times, its diameter provides a limit to the spatial resolution that is obtainable by any method that uses only delay times to determine the model (9, 18).

FWI however is not limited in this way. It seeks to match waveforms, and these are sensitive to sub-Fresnel-zone structure. In this case, it is the wavelength, not the Fresnel zone that is significant (14, 19). Energy scattered from two objects that are separated by more than half a wavelength are distinguishable in terms of their waveforms, in at least some directions, from energy scattered from a single composite object. Provided only that the angular coverage of the target region is sufficiently wide, the spatial resolution of transmission tomographic FWI is of the order of half the incident wavelength. In practice, sufficient angular coverage can be obtained either by locating sources and receivers at all azimuths, or by making use of multiple scattering in a highly heterogeneous medium. In the present context, FWI takes advantage of both approaches.

For neuro imaging as discussed here, the characteristic sound speed is about 1500 m/s, and the maximum frequency employed is about 850 kHz, having a wavelength of about 1.8 mm in the brain. The separation of source and receiver required for transcranial tomography is about 200 mm, giving a Fresnel-zone diameter of about 19 mm. This is the resolution to be expected from conventional time-delay ultrasound computer tomography at 850 kHz, and is the reason that such tomography is not useful for brain imaging. The corresponding resolution for FWI is half the wavelength, that is about 0.9 mm. This is about 20 times better resolved in linear dimension, and about 8,000 times better resolved volumetrically. It is the reason that FWI succeeds to image the brain while time-of-flight tomography cannot.

For conventional time-of-flight tomography to reach the resolution of FWI, time-of-flight tomography would need to use signals that have a Fresnel-zone diameter of about 1 mm. Such signals would require frequencies of several hundred MHz, and these lie far above the bandwidth of signals than can cross the head.

## ACKNOWLEDGMENTS

We gratefully acknowledge assistance in the laboratory from Javier Cudeiro, Thomas Robins and Carlos Cueto. Adrian Umpleby was the lead programmer in the project that developed the Fullwave3D package used in this study.

## SUPPLEMENTARY MEDIA

**Movie 1. Synthesized wavefield crossing the head.** The wavefield from Fig. 1, generated by a single emitter, is shown in time as it propagates across the head.

**Movie 2. Transverse planes.** The target, starting and recovered models from Fig. 3 are shown in three dimensions using a sequence of transverse slices.

**Movie 3. Sagittal planes.** The target, starting and recovered models from Fig. 3 are shown in three dimensions using a sequence of sagittal slices.

**Movie 4. Coronal planes.** The target, starting and recovered models from Fig. 3 are shown in three dimensions using a sequence of coronal slices.

**Sup. Table 1.** Physical properties. Tissue types in the MIDA model [41] together with their corresponding acoustic wave speeds, densities and anelastic absorptions (expressed as quality factor *Q*) used to generate the true model, taken from [21–24,31–36]. The properties of most minor tissue types are estimates. Some properties of major tissue types are not well characterized by available experimental measurements; values of anelastic absorption are especially difficult to measure in the laboratory. Provided that these estimates are broadly correct, their exact values do not influence the quality of the recovered images.

| MIDA number | speed (m/s) | density (kg/m<sup>3</sup>) | quality factor | tissue type |
| --- | --- | --- | --- | --- |
| 1 | 1600.0 | 1174.0 | 156 | Dura |
| 2 | 1511.0 | 1045.0 | 523 | Cerebellum Grey Matter |
| 3 | 1510.0 | 1053.0 | 135 | Pineal Body |
| 4 | 1505.0 | 1044.0 | 1745 | Amygdala |
| 5 | 1515.0 | 1044.5 | 1745 | Hippocampus |
| 6 | 1504.5 | 1007.0 | 20882 | CSF Ventricles |
| 7 | 1510.0 | 1044.5 | 1745 | Caudate Nucleus |
| 8 | 1505.0 | 1044.5 | 1745 | Putamen |
| 9 | 1552.5 | 1041.5 | 302 | Cerebellum White Matter |
| 10 | 1505.0 | 1044.5 | 1745 | Brain Grey Matter |
| 11 | 1546.3 | 1045.5 | 299 | Brainstem Midbrain |
| 12 | 1552.5 | 1041.0 | 302 | Brain White Matter |
| 13 | 1542.0 | 1075.0 | 30 | Spinal Cord |
| 14 | 1546.3 | 1045.5 | 299 | Brainstem Pons |
| 15 | 1546.3 | 1045.5 | 176 | Brainstem Medulla |
| 16 | 1505.0 | 1044.5 | 1745 | Nucleus Accumbens |
| 17 | 1505.0 | 1044.5 | 1745 | Globus Pallidus |
| 18 | 1629.5 | 1075.0 | 146 | Optic Tract |
| 19 | 1515.0 | 1053.0 | 135 | Hypophysis or Pituitary Gland |
| 20 | 1505.0 | 1044.5 | 1745 | Mammillary Body |
| 21 | 1520.0 | 1044.5 | 1745 | Hypothalamus |
| 22 | 1552.5 | 1041.0 | 302 | Commissura (Anterior) |
| 23 | 1552.5 | 1041.0 | 302 | Commissura (Posterior) |
| 24 | 1578.2 | 1049.8 | 841 | Blood Arteries |
| 25 | 1578.2 | 1049.8 | 841 | Blood Veins |
| 26 | 343.0 | 1.2 | 233990 | Air Internal - Ethmoidal Sinus |
| 27 | 343.0 | 1.2 | 233990 | Air Internal - Frontal Sinus |
| 28 | 343.0 | 1.2 | 233990 | Air Internal - Maxillary Sinus |
| 29 | 343.0 | 1.2 | 233990 | Air Internal - Sphenoidal Sinus |
| 30 | 343.0 | 1.2 | 233990 | Air Internal - Mastoid |
| 31 | 343.0 | 1.2 | 233990 | Air Internal - Nasal/Pharynx |
| 32 | 1504.5 | 1007.0 | 20882 | CSF General |
| 33 | 1504.5 | 1007.0 | 20882 | Ear Cochlea |
| 34 | 1504.5 | 1007.0 | 20882 | Ear Semi-circular Canals |
| 35 | 1639.6 | 1099.5 | 4358 | Ear Auricular Cartilage (Pinna) |
| 36 | 3514.9 | 1908.0 | 16 | Mandible |
| 37 | 1620.7 | 1102.0 | 16 | Mucosa |
| 38 | 1588.4 | 1090.4 | 278 | Muscle (General) |
| 39 | 1639.6 | 1099.5 | 4358 | Nasal Septum (Cartilage) |
| 40 | 2813.7 | 1908.0 | 20 | Skull (General) |
| 41 | 4565.9 | 2180.0 | 11 | Teeth |
| 42 | 1588.4 | 1090.4 | 682 | Tongue |
| 43 | 1440.2 | 911.0 | 501 | Adipose Tissue |
| 44 | 3514.9 | 1908.0 | 16 | Vertebra - C1 (atlas) |
| 45 | 3514.9 | 1908.0 | 16 | Vertebra - C2 (axis) |
| 46 | 3514.9 | 1908.0 | 16 | Vertebra - C3 |
| 47 | 3514.9 | 1908.0 | 16 | Vertebra - C4 |
| 48 | 3514.9 | 1908.0 | 16 | Vertebra - C5 |
| 49 | 1639.6 | 1099.5 | 4358 | Intervertebral Discs |
| 50 | 1480.0 | 1000.0 | 20000 | Background |
| 51 | 1624.0 | 1109.0 | 91 | Epidermis/Dermis |
| 52 | 2117.5 | 1178.3 | 32 | Skull - Diploë |
| 53 | 2813.7 | 1908.0 | 20 | Skull - Inner Table |
| 54 | 2813.7 | 1908.0 | 20 | Skull - Outer Table |
| 55 | 1643.3 | 1075.5 | 2868 | Eye Lens |
| 56 | 1638.8 | 1032.0 | 1633 | Eye Retina/Choroid/Sclera |
| 57 | 1525.8 | 1004.5 | 412 | Eye Vitreous |
| 58 | 1562.5 | 1050.5 | 5464 | Eye Cornea |
| 59 | 1525.8 | 1004.5 | 412 | Eye Aqueous |
| 60 | 1588.4 | 1090.4 | 278 | Muscle - Platysma |
| 61 | 1750.0 | 1142.0 | 124 | Tendon - Galea Aponeurotica |
| 62 | 1477.0 | 911.0 | 304 | Subcutaneous Adipose Tissue |
| 63 | 1588.4 | 1090.4 | 278 | Muscle - Temporalis/Temporoparietalis |
| 64 | 1588.4 | 1090.4 | 278 | Muscle - Occipitiofrontalis - Frontal |
| 65 | 1588.4 | 1090.4 | 278 | Muscle - Lateral Pterygoid |
| 66 | 1588.4 | 1090.4 | 278 | Muscle - Masseter |
| 67 | 1588.4 | 1090.4 | 278 | Muscle - Splenius Capitis |
| 68 | 1588.4 | 1090.4 | 278 | Muscle - Sternocleidomastoid |
| 69 | 1588.4 | 1090.4 | 278 | Muscle - Occipitiofrontalis - Occipital |
| 70 | 1588.4 | 1090.4 | 278 | Muscle - Trapezius |
| 71 | 1588.4 | 1090.4 | 278 | Muscle - Mentalis |
| 72 | 1588.4 | 1090.4 | 278 | Muscle - Depressor Anguli Oris |
| 73 | 1588.4 | 1090.4 | 278 | Muscle - Depressor Labii |
| 74 | 1588.4 | 1090.4 | 278 | Muscle - Nasalis |
| 75 | 1588.4 | 1090.4 | 278 | Muscle - Orbicularis Oris |
| 76 | 1588.4 | 1090.4 | 278 | Muscles - Procerus |
| 77 | 1588.4 | 1090.4 | 278 | Muscle - Levator Labii Superioris |
| 78 | 1588.4 | 1090.4 | 278 | Muscle - Zygomaticus Major |
| 79 | 1588.4 | 1090.4 | 278 | Muscle - Orbicularis Oculi |
| 80 | 1588.4 | 1090.4 | 278 | Muscle - Levator Scapulae |
| 81 | 1588.4 | 1090.4 | 278 | Muscle - Medial Pterygoid |
| 82 | 1588.4 | 1090.4 | 278 | Muscle - Zygomaticus Minor |
| 83 | 1588.4 | 1090.4 | 278 | Muscles - Risorius |
| 84 | 1588.4 | 1090.4 | 278 | Muscle - Buccinator |
| 85 | 343.0 | 1.2 | 233990 | Ear Auditory Canal |
| 86 | 343.0 | 1.2 | 233990 | Ear Pharyngotympanic Tube |
| 87 | 2117.5 | 1178.3 | 32 | Hyoid Bone |
| 88 | 1559.5 | 1048.0 | 216 | Submandibular Gland |
| 89 | 1559.5 | 1048.0 | 216 | Parotid Gland |
| 90 | 1559.5 | 1048.0 | 216 | Sublingual Gland |
| 91 | 1588.4 | 1090.4 | 278 | Muscle - Superior Rectus |
| 92 | 1588.4 | 1090.4 | 278 | Muscle - Medial Rectus |
| 93 | 1588.4 | 1090.4 | 278 | Muscle - Lateral Rectus |
| 94 | 1588.4 | 1090.4 | 278 | Muscle - Inferior Rectus |
| 95 | 1588.4 | 1090.4 | 278 | Muscle - Superior Oblique |
| 96 | 1588.4 | 1090.4 | 278 | Muscle - Inferior Oblique |
| 97 | 343.0 | 1.2 | 233990 | Air Internal - Oral Cavity |
| 98 | 1750.0 | 1142.0 | 124 | Tendon - Temporalis Tendon |
| 99 | 1505.0 | 1044.5 | 1745 | Substantia Nigra |
| 100 | 1552.5 | 1041.0 | 302 | Cerebral Peduncles |
| 101 | 1629.5 | 1075.0 | 146 | Optic Chiasm |
| 102 | 1629.5 | 1075.0 | 146 | Cranial Nerve I - Olfactory |
| 103 | 1629.5 | 1075.0 | 146 | Cranial Nerve II - Optic |
| 104 | 1629.5 | 1075.0 | 146 | Cranial Nerve III - Oculomotor |
| 105 | 1629.5 | 1075.0 | 146 | Cranial Nerve IV - Trochlear |
| 106 | 1629.5 | 1075.0 | 146 | Cranial Nerve V - Trigeminal |
| 107 | 1629.5 | 1075.0 | 146 | Cranial Nerve V2 - Maxillary Division |
| 108 | 1629.5 | 1075.0 | 146 | Cranial Nerve V3 - Mandibular Division |
| 109 | 1629.5 | 1075.0 | 146 | Cranial Nerve VI - Abducens |
| 110 | 1629.5 | 1075.0 | 146 | Cranial Nerve VII - Facial |
| 111 | 1629.5 | 1075.0 | 146 | Cranial Nerve VIII - Vestibulocochlear |
| 112 | 1629.5 | 1075.0 | 146 | Cranial Nerve IX - Glossopharyngeal |
| 113 | 1629.5 | 1075.0 | 146 | Cranial Nerve X - Vagus |
| 114 | 1629.5 | 1075.0 | 146 | Cranial Nerve XI - Accessory |
| 115 | 1629.5 | 1075.0 | 146 | Cranial Nerve XII - Hypoglossal |
| 116 | 1505.0 | 1044.5 | 1745 | Thalamus |

